# Mechanisms for action prediction operate differently in observers with motor experience

**DOI:** 10.1101/044438

**Authors:** Lincoln J Colling, William F Thompson, John Sutton

## Abstract

Recent theoretical and empirical work has suggested an important role for the motor system in generating predictions about the timing of external events. We tested the hypothesis that motor experience with an observed action changes how observers generated predictions about these actions by comparing the performance of naïve and experienced observers on a task that required participants to predict the timing of particular critical points in an ongoing observed action. Crucially, we employed action and non-action stimuli with identical temporal dynamics, and we predicted that motor experience would enhance prediction accuracy specifically for actions and would have a reduced or negligible effect on enhancing prediction accuracy for non-action stimuli. Our results showed that motor experience did modulate prediction accuracy for action stimuli relative to non-action stimuli. No difference between conditions was observed for the naïve observers.

## Introduction

Joint action relies crucially on the ability to predict the actions of other agents. In many cases of joint action, it is important that people can predict not only *what* action their co-actor will preform but also *when* they will perform it. Indeed, in the case of joint actions such as music and dance performance, ones co-actor’s actions can be known in advance. In these cases, the crucial predictions are about the *timing* of ones co-actor’s actions. The important role of prediction has been emphasised in many recent theoretical accounts of joint action coordination (Colling and Williamson 2014; Csibra 2008; Wilson and Knoblich 2005; Knoblich, Butterfill, and Sebanz 2011; Colling, Knoblich, and Sebanz 2013). Furthermore, as many cases of joint action, such as music, dance, or sports performance, also involve cases of *expert* performance, researchers have also turned their attention to how these predictive processes might be modulated by experience. Much of this work, however, has focused on paradigms where people are required to predict action outcomes—that is, *what* the co-actor will do (e.g., Sebanz and Shiffrar 2009; Aglioti, Cesari, Romani, and Urgesi 2008). Even in paradigms where speeded judgements are required (e.g., Urgesi, Savonitto, Fabbro, and Aglioti 2012) questions of how observers might be able to generate predictions about the *timing of observed actions* have tended to be neglected. Therefore, it is the influence of first-hand action experience on the prediction of action timing that is the primary concern of the present study.

### Neural mechanisms for action observation and prediction

There is a rich literature examining the influence of motor experience on neural networks that are preferentially activated during action observation (for seminal examples, see Calvo-Merino, Glaser, Grézes, Passingham, and Haggard 2005; Calvo-Merino, Grézes, Glaser, Passingham, and Haggard 2006; Cross, Hamilton, and Grafton 2006). The initial work on this action observation network began with the discovery of *mirror neurons* in premotor regions of the monkey brain. Mirror neurons are active when the monkey performs an action and also when the monkey observes the same, or a similar, action performed by another (Rizzolatti, Fadiga, Gallese, and Fogassi 1996; Gallese, Fadiga, Fogassi, and Rizzolatti 1996; and for a recent review see Giese and Rizzolatti 2015). Many theories about the functional role of the mirror system have focused on how the mirror system might play a role in action recognition or action understanding (e.g., Sartori and Betti 2015; Giese and Rizzolatti 2015). Other accounts, however, suggest that this system might also, or even primarily, be involved in generating predictions about ongoing observed actions (e.g., Wilson and Knoblich 2005; Csibra 2008; Colling et al. 2013; and see Kilner, Friston, and Frith 2007, for an account of the role of the mirror system in action prediction in the context of action understanding).

Accounts linking the mirror system to action prediction have relied on the fact that the mirror system is partially co-extensive with the action control system and, therefore, observed actions might be processed by some of the same neural machinery involved in planning and executing actions. This link to planning and executing actions is important, because work on computational models of action control highlight a fundamental role for *prediction* in action control (Wolpert 1997).

### Computational models of action control

Concepts borrowed from control theory have been particularly useful for understanding how prediction, during both action observation and action control, might be achieved (for an introduction to control theory, see Golnaraghi 2010). Specifically, *inverse models* and *forward models* have proven theoretically useful. Inverse models perform an *inverse mapping* from an output or goal state to the sequence of control commands necessary to produce that output.

And forward models perform a *forward mapping* from the control commands to the output. That is, they model the dynamics of the system being controlled.

Inverse and forward models—together known as *internal models—*have a central role in theoretical accounts of action control (for example, see Wolpert, Miall, and Kawato 1998).

Inverse models act as controllers that transform the desired limb trajectory into the motor commands that would produce that trajectory. And forward models replicate the dynamics of the limb and can, therefore, be used to predict how the limb will respond to motor commands (Wolpert and Kawato 1998). Running the forward model *offline—*that is, without producing any actual motor output—can be used to internally simulate limb movements. Grush (1997; 2004) refers to this process as *emulation* and to the forward model as an *emulator*.

### Predicting observed actions

Grush’s (1997; 2004) ideas about emulation have been developed into an account of action prediction that has been termed the *emulator theory of action prediction* (Colling and Williamson 2014; Colling, Thompson, and Sutton 2014). While many slightly varying formulations exist (see also Colling et al. 2013; Csibra 2008; Keller 2012; Wilson and Knoblich 2005; Wolpert, Doya, and Kawato 2003; Vesper, Butterfill, Knoblich, and Sebanz 2010), the basic idea is that prediction of observed actions relies on the same internal mechanisms that support action production. The basic claim is that the observer’s action control system acts as an emulator enabling the observed action to be internally simulated in real-time. These real-time simulations can then be used as the basis for anticipatory action planning. However, in order to internally simulate the observed action using an emulator, a motor command, which ordinarily drives the forward model during action production, is needed. One way to generate this motor command might be to formulate a conjecture about what action the observed agent is producing (Kilner et al. 2007) or by visual analysis of the observed action (Csibra 2008). Visual analysis can be coupled with an inverse model to simulate the motor commands driving the observed action.

A key claim of the emulator theory of action prediction, at least as formulated by Wilson and Knoblich (2005) and Colling and colleagues (Colling et al. 2013; Colling and Williamson 2014), is that the observed action is mapped onto the observer’s body in a part-by-part manner. That is, prediction occurs by internally simulating the action *as if the observer was performing it*. Because prediction is tied to the observer’s own action control system, predictions should carry traces of this system.

One way to look for these traces is to examine the influence of specific motor expertise on action prediction. One such example comes from Aglioti et al. (2008). This study employed a basketball free throw prediction task to compare the performance of novice and expert basketball players. The general finding from these paradigms is that experts generate more accurate predictions than novices (Abernethy 1990; Isaacs and Finch 1983; Aglioti et al. 2008; Sebanz and Shiffrar 2009). Although studies comparing action prediction in experts and novices appear to demonstrate that predictive processes are enhanced by motor experience at least one concern can be raised. Specifically, the causal relationship between expertise and prediction is not clear. It may be the case that expertise causes superior predictive abilities; however, it is also possible that those who become experts do so because they already possess superior predictive abilities. To uncover the direction of causality it may be preferable to train people on an action rather than use experts.

In a study by Urgesi et al. (2012, Experiment 2) this is indeed the approach that was adopted. In this experiment, participants were given different types of volleyball training that were either general or specific to the volleyball prediction task that they would later perform. The results showed that training led to enhanced prediction abilities on a volleyball shot prediction task (predicting whether a volleyball shot would land in or out of the court) with observers making quicker and more accurate judgements.

While studies such as these have been able to demonstrate that experience with actions can lead to enhanced prediction abilities, and even the ability to generate these predictions quicker, they neglect to address the question that is central to the current work. Specifically, these paradigms all involve tasks where participants are required to predict action outcomes. That is, participants are required to predict *what* action the observed agent will perform.

Consequently, these paradigms do not provide a measure of temporal prediction accuracy. That is, they don’t require participants to generate a prediction about the temporal properties of the action, or about *when* the action will be performed.

Indeed, most previous work by, for example, Aglioti et al. (2008), Sebanz and Shiffrar (2009), Ikegami and Ganesh (2014), Mulligan and Hodges (2013), and others^1^, participants were asked to generate a prediction about the outcome of an action. For example, whether a basketball free-throw would be successful or not. While these tasks do test predictive mechanisms, it is not clear whether they test the same predictive mechanisms that underlie joint performance in music, dance, and sport. The predictive mechanisms that underlie joint action must have two features, neither of which are tested by these kinds of tasks. First, predictions must be about the temporal properties of the observed action—that is, it isn’t sufficient to simply predict what action will be performed or the outcome of an action rather the key lies in predicting *when* the action will be performed. And second, it must be possible to use these predictions as the basis for anticipatory action planning. This second concern is highlighted by recent work from Mann, Abernethy, and Farrow (2010). In this study, participants were required to generate a prediction about an action and then report their prediction in different ways. Responses could be made through a verbal report or by producing the appropriate action in response to the prediction (in this case, performing the correct cricket shot in response to the predicted trajectory of a ball delivered by a bowler). The results showed that the accuracy of predictions was modulated by response modality, suggesting that predictions generated for a verbal report and action planning might reside in different (sub)systems. Therefore, to examine the action prediction mechanisms that might underlie joint action, it is necessary to employ tasks that replicate (at least some) of the coordination demands found in joint action. One example of this can be found in temporal alignment tasks.

### Temporal alignment tasks

Temporal alignment tasks are tasks requiring observers to perform and action on the basis of their prediction, rather than make a decision. Asking participants to perform an action on the basis of a temporal prediction was the approach employed by Cross, Stadler, Parkinson, Schütz-Bosbach, and Prinz (2013). In this study, participants were asked to generate a prediction about when a gymnast or a toy, which was moving across the screen, would reappear after moving behind an occluder. Participants were required to press a button at the point in time that they believed the person or toy would reappear. The primary finding of this study was that repeated visual exposure to the stimuli resulted in more accurate predictions about when the gymnast or toy would reappear.

Importantly, however, it is not clear whether this task actually taps into action prediction mechanisms. Rather, this task could be performed using mechanisms that allow people to judge the duration of intervals. As the gymnast or toy moves across the screen, accurate perception of how long it takes to move a fixed distance would allow the observer to accurately predict when it will reappear from behind the occluder.

A different task, developed by Colling et al. (2014)^2^, was designed to tap into mechanisms specifically related to action prediction, and to test the claim of the emulator hypothesis that traces of the observer’s action control system should be evident in the predictions that they generate. In this temporal alignment task, participants viewed mannequins performing up-and-down arm movements while attempting to align a button press with the apex of each upward movement (when it changed from upward to downward).

Importantly, the spacing between the points of direction change was irregular thus preventing the observer from relying on interval timing mechanisms (see Colling et al. 2014, Experiment 2–3). Mannequins were viewed under two conditions. In the *self* condition, participants viewed mannequins created from motion capture recordings of their own movements produced at an earlier time. In the *other* condition, participants viewed mannequins created from recordings of another person’s movements. The logic of this manipulation was that if people generate predictions using their own action control system, with forward models that replicate their own action dynamics, then predictions in the self condition should be more accurate, because in the self condition the dynamics of the predictor and the dynamics of the predicted action are matched.

The results confirmed this and people were significantly more accurate at aligning button press responses with recordings of their own actions. Importantly, these tasks require participants to not only generate predictions quickly and in real-time but also to plan and execute actions on the basis of these predictions. By employing a paradigm such as this, it should be possible to examine the influence of motor experience on prediction in tasks that more closely match the coordination demands found in joint action.

### How does motor experience change action prediction?

While we have highlighted how previous research has tended to neglect the impact of experience on the prediction of action timing, there is also a more fundamental concern about this work that motivates the present study. The studies cited above (e.g., Aglioti et al. 2008) suggest that motor experience enables more accurate predictions (at least for action outcome tasks); however, these studies do not answer the question of *how* the prediction process changes to achieve this. For instance, it might just be that experts and novices employ the same strategy or mechanisms and that experts are just able to employ this strategy with greater efficiency or accuracy. This might be because experts have more accurate, or even new, sensorimotor mappings that are laid down by extensive practise (see, for example Calvo-Merino et al. 2005; Calvo-Merino et al. 2006; Cross et al. 2006).

However, another way in which motor experience might modulate action prediction is by changing the way in which sensory-motor mappings are deployed—that is, it might be the case that novices and experts employ distinct strategies, with the consequence of this being superior prediction accuracy for the experts^3^. The suggestion that distinct processes or strategies might underlie action prediction in experienced and naïve observers is found in an extension of the emulator hypothesis, developed by Schubotz (2007). Based on results from fMRI (e.g., Schubotz and von Cramon 2004) and lesion studies (e.g., Schubotz, Sakreida, Tittgemeyer, and von Cramon 2004), which implicate premotor regions in sequence prediction, Schubotz (2007) suggests that motor simulation is a general purpose predictive mechanism for predicting not only human actions but all manner of external events. In the case of *reproducible events* (e.g., human actions) it is possible to internally simulate the observed action using the same mechanisms used to produce them, as claimed by the emulator theory (e.g., see Colling et al. 2013; Wilson and Knoblich 2005; Colling and Williamson 2014). In the terminology of Schubotz (2007, p. 213), observers use their “motor memories to run a simulation of the observed movement”. In the absence of these motor memories, Schubotz (2007) suggests that predictions might be generated by mapping the observed event onto an effector that best matches the general dynamics of the observed stimuli. Therefore, it might be that experienced and naïve observers actually utilise distinct strategies during action prediction, with experienced observers internally replicating the observed action *as it was performed* and naïve observers just replicating the stimulus dynamics with whatever effector does the best job. This generic simulation (as opposed to *action specific* simulation) might not only occur in the absence of motor experience. It might also occur when the observed stimuli cannot be easily mapped onto the observer’s body. For example, when the action stimuli are impoverished so that it is not clear *how* the action is being produced—that is, when the observed actions are not amenable to visual analysis (Csibra 2008)—it might not be clear which action, out of all possible actions, to internally simulate. In this case, observers might again internally replicate the dynamics of the stimulus using whatever effector does the best job rather than simulating the actual action.

### Aims of the present study

The aim of the present study is to investigate *how* predictive processes change when observers have experience producing observed actions. Previous studies have reported that action prediction becomes more accurate when observers possess motor experience; however, it is not clear how the process changes to enable this. Indeed, measuring prediction accuracy alone may not be sufficient to do this. Furthermore, previous studies on action prediction have generally tended to focus on predicting action outcomes, with relatively few studies (e.g., Colling et al. 2014; Keller et al. 2007; Flach et al. 2003) employing the kind of tasks that replicate the temporal demands found in joint action.

The work of Schubotz (2007) suggests that experienced and naïve observers might employ distinct mechanisms for action prediction. It might be possible to test whether experienced and naïve observers engage distinct predictive mechanisms, or employ different predictive strategies, by designing a manipulation that should affect only one strategy and not the other. Schubotz (2007) suggests that generic, non-action specific, simulation should occur when observers have little or no experience producing the observed action. Furthermore, action-specific simulation should also be difficult when the stimulus is impoverished—for example when it does not clearly depict an action—so that is cannot easily be mapped on to the body in a part-by-part manner. Furthermore, we might expect these two factors to interact. That is, if an observer is engaged in generic, or approximate, simulation of the observed action, such as when they have no experience producing the action, then diminishing stimulus detail, so that the stimulus cannot be mapped onto the body, should be of little consequence, because generic simulation does not require the stimulus to be mapped onto the body.

However, if the observer is engaged in a detailed part-by-part simulation then decreasing the stimulus detail so that it is not clear how the action is being produced, should disrupt the predictive process. That is, changing the nature of the stimulus, so that action-specific information—that is, information about how the action is performed—but not the critical dynamic information is reduced, should only affect the action specific strategy.

To test this hypothesis, we examined the influence of motor experience on prediction during a temporal alignment task similar to that used by Colling et al. (2014). Two groups of participants, those with experience producing the observed action and naïve participants, viewed actions under two conditions. In the full information condition, the stimuli depicted the actions in full, including information about the arrangement of the limbs and joints during the production of the action. In the point information condition, participants were required to align a response with the same dynamic information; however, the displays were impoverished so that they did not depict an action. We predict that first hand experience producing the action will lead to a switch in prediction strategy such that for experienced observers decreasing stimulus information should hamper the process of internal replication while naïve observers, who only engage an approximate, rather than an action specific, predictive solution should display little, or no, difference between the two conditions. This should result in a difference in prediction accuracy between stimulus conditions as a function of motor experience—that is stimulus condition and motor experience should interact. Using this procedure has an advantage over simply comparing prediction accuracy for a single stimulus type (e.g., as typically done in studies of sports expertise such as, for example, Aglioti et al. 2008) because the stimulus manipulation is predicted to have a different effect on alignment accuracy depending on the strategy employed by the observer. Thus it may be possible to examine whether the naïve and experienced group employ distinct strategies.

Typical sports expertise studies are blind to whether experts employ a different, more accurate, strategy relative to novices or whether experts just employ the same strategy as novices but with greater precision. Examining the interaction also has an additional benefit in that it might be the case that added visual complexity makes the stimulus easier to visually track. However, if this is the case, then we can expect to see the same stimulus-related advantage in prediction accuracy regardless of motor experience.

## Methods

### Participants

There were 13 participants (11 females, mean age of 27.5 years) in the experienced group, and 12 participants (8 females, mean age of 20.1 years)^4^ in the naïve group. All participants were right-handed (Oldfield 1971), and all procedures were approved by the Macquarie University Human Subjects Ethics committee.

### Stimuli

To create the stimuli for the test session, five right-handed females (mean age of 24.8 years) performed a movement task while their movements were recorded with motion capture. The movement task involved tracing out wave and zigzag patterns as if drawing them on a blackboard (the patterns measured 0.584 m × 0.841 m; see Figure 1). Each pattern contained five peaks, alternating in height from large to small; however, they differed in the nature of the direction change at the apex of the peaks. The zigzag pattern contained an abrupt change while the wave pattern had a smooth direction change. The two patterns were only used to keep the actors engaged in the task by providing some variety, and we had no predictions about how the pattern would influence performance; therefore, the data were collapsed over pattern during data analysis^5^.

**Figure 1.**
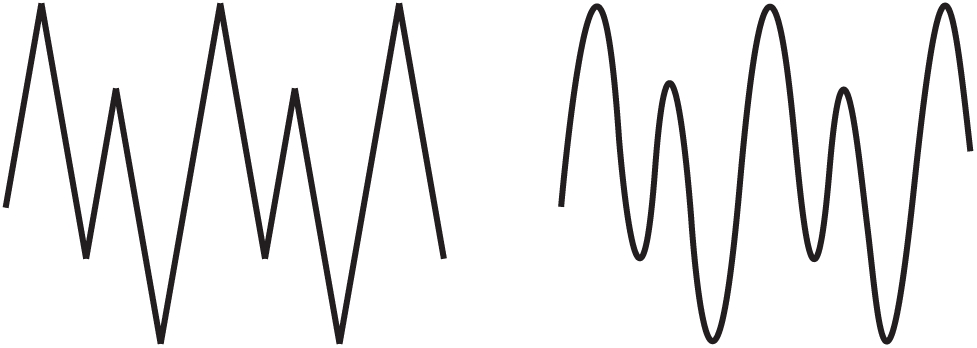
The zigzag (left) and wave (right) patterns used as stimuli during the movement task

Movements were recorded using an 8-camera 3-D passive optical motion capture system (Vicon MX with 4 MX-F20 and 4 MX13+ cameras; 200 Hz sampling rate). To define the limb segments and the position of the torso, markers were placed on the shoulders, the right elbow, wrist, waist, and the top of the right hand (See Figure 2). For the full information condition, the motion capture data was rendered as an animated character consisting of an upper torso, right arm and right hand. For the point information condition, only a single point tracking the movement of the RFIN marker was displayed (See Figure 3). Mannequins were preferred over point-light displays because they preserve occlusion.

**Figure 2.**
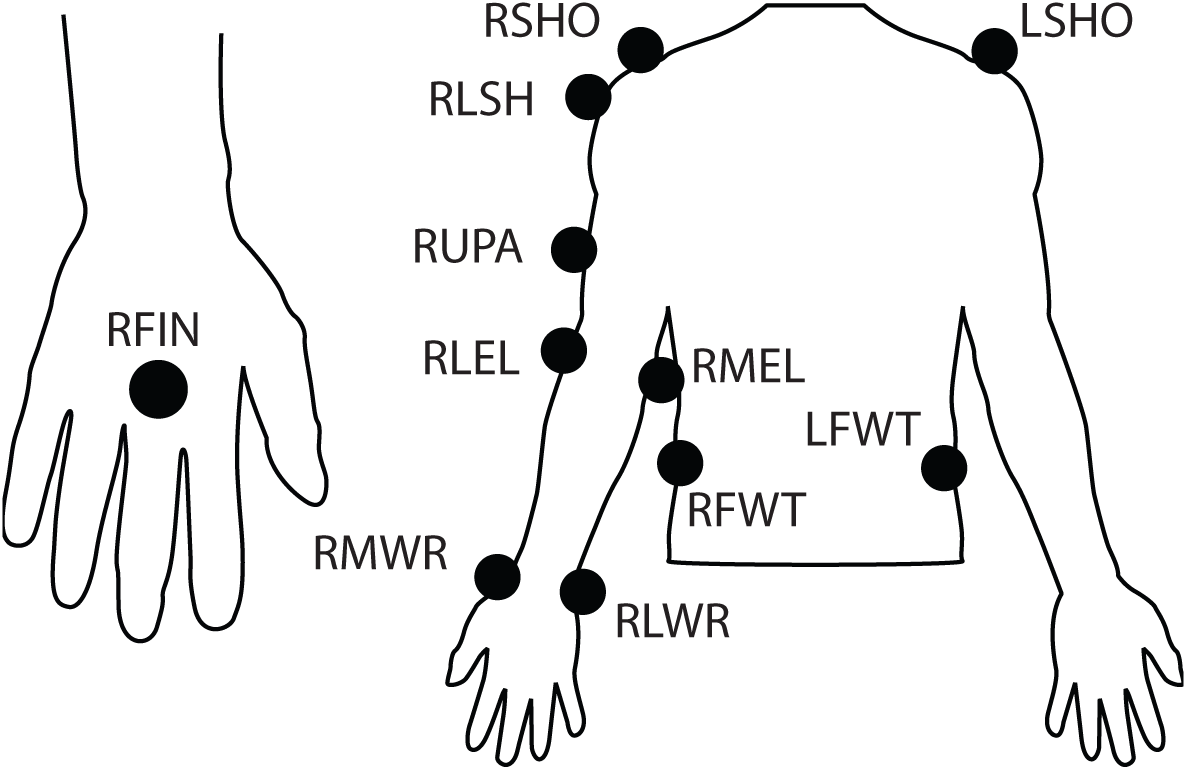
Marker positions for the 11 reflective markers used during the movement task

**Figure 3.**
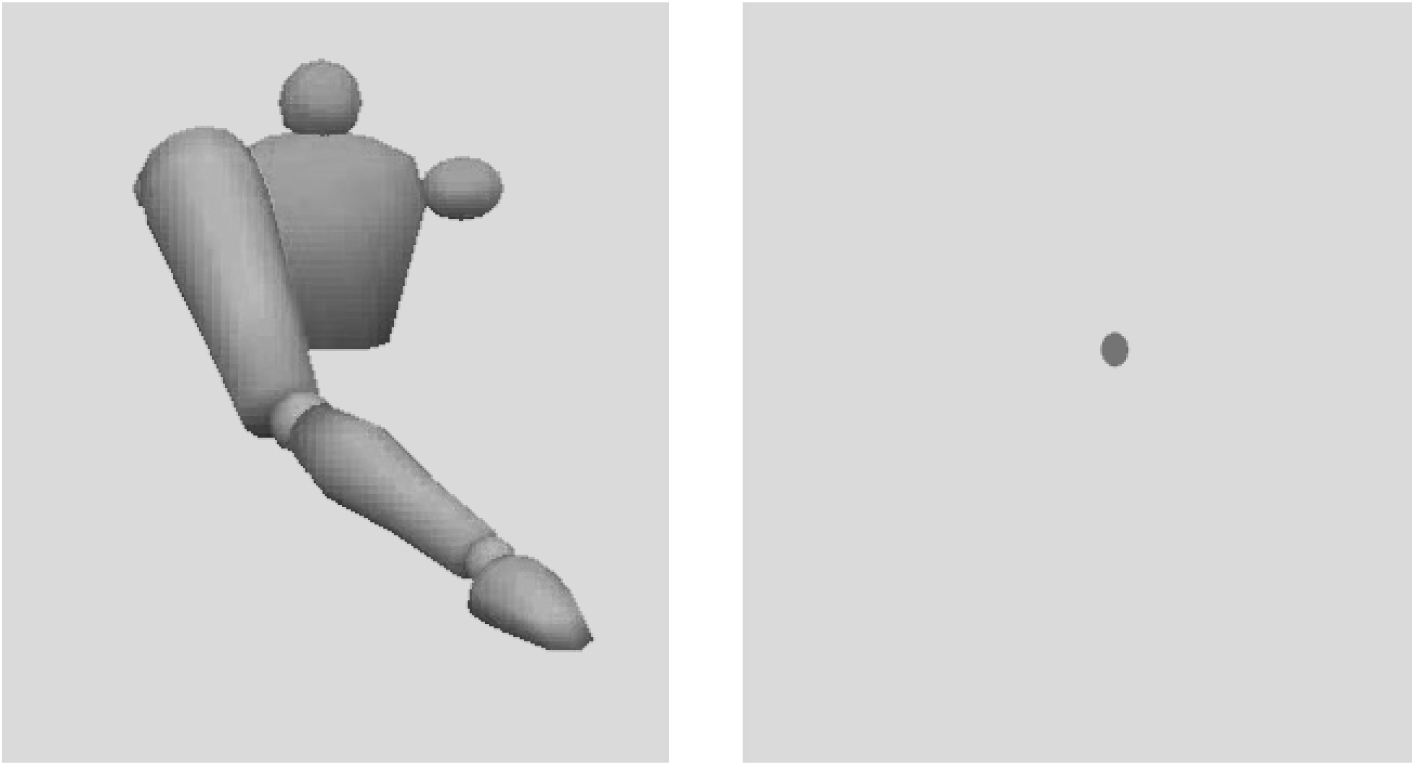
Example stimuli from the full information condition (left) and the point information condition (right)

### Procedure

Participants in the experienced group undertook a movement session that was identical to the task employed during stimulus creation. Participants performed 3 blocks containing 5 repetitions of each pattern (in random order) with their eyes closed to limit visual experience. The movement session and the test session were on average separated by 16.69 days (7 to 28 days).

The task in the test session, which was conducted in a different lab to the movement task, was to press the response button when the hand of the mannequin, or the marker tracking the hand, reached the apex of each upward movement. That is, on each trial participants were required to press the button *five* times. They were instructed to synchronise the button-press with the display as accurately as possible and were told that this may require them to anticipate when the peak will occur. Each participant performed 4 blocks containing 40 unique stimuli (composed of 20 trials for the full information condition and 20 trials for the point information condition) with equal numbers of wave and zigzag stimuli. Participants that did not undergo the movement session were given a brief verbal description of the movement task; however, the task instructions were identical across the two groups.

### Statistical analyses

To measure alignment accuracy, we calculated the absolute timing difference between the occurrence of the peak in the motion capture trajectory and the occurrence of the button presses performed by the participant. Only the last four button presses were analysed because several stimuli contained missing frames leading up to the first peak. The primary analysis of interest was whether the difference between the full information condition and the point information condition was modulated by group membership. To examine this, we first calculated what we termed the *full information advantage*. The full information advantage is simply the difference in the timing error between the full information condition and the point information condition. We hypothesised that the full information advantage would be larger in the motor experience group relative to the naïve group.

In addition to examining the full information advantage we also examined the difference in absolute timing error between the experienced group and the naïve group when the data was averaged across the two condition and we examined the difference in absolute timing error between the point information condition and the full information condition when the data were not separated by group. This was done to examine whether there was an overall difference between the two conditions that was independent of group or between the two groups that was independent of condition.

To perform these comparisons, we adopted the approach outlined by Kruschke (2013), which involves fitting a *t*-distribution to the data by estimating the three parameters of the *t*-distribution: a mean (*µ*), a standard deviation (*σ*), and a shape parameter (*ν*)—the addition of the shape parameter (*ν*) allows the model to account for outliers in the data^6^. This produces a full posterior distribution of credible values for each parameter. For independent samples analyses (that is, the two between-group comparisons) independent distributions are fit to the data and the difference is examined by subtracting the posteriors. For dependent samples analyses (that is, the between condition comparison and the two posthoc comparisons) paired differences are first calculated and then the distribution is fit to the difference scores. We report the mean as well as the 95% highest density intervals (HDI) for the parameter *µ*, as well as the 95% HDI for the effect size estimates, designated as *d*, which are calculated as *µ/σ* for the dependent sample comparisons and as 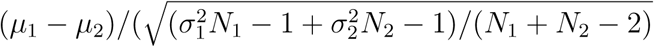 for independent samples comparisons. Highest density intervals can be interpreted as indicating the bounds that contain 95% of credible values for the parameter. The width of the HDIs provides a measure of the precision of the estimates. All the raw data and analysis code necessary to reproduce the results and the figures is available online (see http://osf.io/ebf45).

## Results

The descriptive statistics are shown in Table 1. The table shows sample mean and standard deviation for each condition (in each group and collapsed across group) together with the correlations between the conditions. The sample means and standard deviations for each group with the data collapsed across condition are also shown. The results of the parameter estimates showed that, when the data was collapsed across stimulus condition, there were no systematic differences in mean alignment accuracy between the experienced group (posterior mean = 123.94 ms, 95% HDI[92.82; 153.54]) and the naïve group (posterior mean = 112.99 ms, 95% HDI[90.84; 134.37]), posterior mean difference = 10.94 ms, 95% HDI[-25.35; 48.37], *d* = 0.28, 95% HDI[−0.59; 1.05]. The mean group difference, and 95% HDIs, are shown in Figure 4A. Furthermore, when the data was collapsed across group, there were no systematic differences in the alignment accuracy between the point information condition (posterior mean = 119.15 ms, 95% HDI[102.71; 136.7]) and the full information condition (posterior mean = 117.27 ms, 95% HDI[99.64; 134.82]) posterior mean difference = 2.22 ms, 95% HDI[−1.42; 6.04], *d* = 0.25, 95% HDI[−0.16; 0.67]. The mean condition difference, and 95% HDI, are shown in Figure 4B.

**Figure 4.**
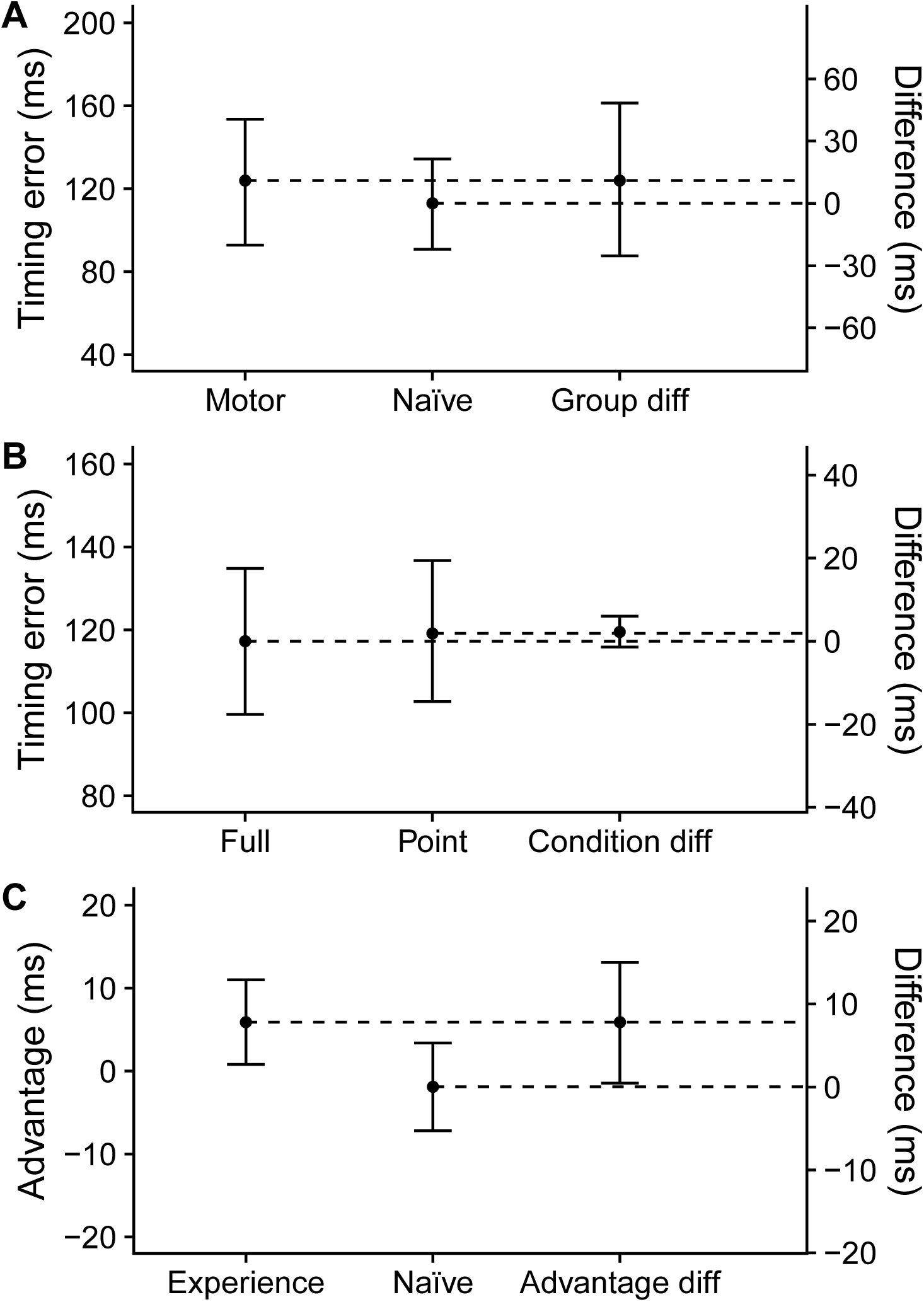
(A) Plot of the Group main effect. The plot shows the mean of the posterior and the 95% HDI of each group as well as the posterior difference between the two groups. (B) Plot of the Condition main effect. The plot shows the mean of the posterior and the 95% HDI of each condition as well as the posterior difference between the two conditions. (C) Plot of the full information advantage. The plot shows the mean of the posterior and the 95% HDI for each group as well as the posterior difference between the two groups.

**Table 1.** Sample mean and sample standard deviation for each condition in each group and combined across groups, together with the sample mean and sample standard deviation for each group combined across condition. The correlations between conditions, at the level of the group and when combined across groups, is also shown.

| Condition | Experienced (n = 13) | Naïve (n = 12) | Combined (n = 25) |
| --- | --- | --- | --- |
| Point | 128.11 (46.48) | 112.4 (29.64) | 120.57 (39.33) |
| Full | 121.98 (46.74) | 114.13 (35.67) | 118.21 (41.13) |
| Combined | 125.04 (46.44) | 113.26 (32.57) | — |
| Correlation | 0.99 | 0.99 | 0.98 |
| Difference | 6.13 (8.04) | -1.74 (7.77) | — |

Our primary comparison of interest, however, was whether the effect of stimulus condition was modulated by group assignment. To test, we compared the magnitude of the *full information advantage*—that is, the magnitude of *difference* between the the full information condition and the point information condition—between the two groups. The results of this analysis revealed reliable differences in the size of the full information advantage between the experienced group and the naïve group, posterior mean difference = 7.79 ms, 95% HDI[0.43; 15], *d* = 0.91, 95% HDI[0.04; 1.8]. Examining the size of the full information advantage separately in each group separately revealed a reliable advantage for full information condition in the motor experience group (posterior mean = 5.89 ms, 95% HDI[0.8; 11.01], *d* = 0.25, 95% HID[0.07; 1.37]); however, in the naïve group, credible values for this difference were both positively and negatively signed indicating no reliable difference (posterior mean = −1.9 ms, 95% HDI[-7.2; 3.39], *d* = −0.22, 95% HID[−0.87; 0.36]). The full information advantage difference is shown in Figure 4C.

### Exploratory analysis of group differences

A further attempt was made to explore differences in task performance between the experienced and naïve group. To do this, we examined whether there were any differences in task performance related to whether participants primarily responded to the local or the global dynamics of the stimulus. In the stimulus, the duration of each upward movement alternated from long to short leading to local variations in the timing of the peaks. That is, the timing of the peaks was not evenly spaced across the trial with peaks being separated by alternating long and short gaps. If participants based their responses on the global dynamics of the stimulus—for example, the average inter-peak interval—and produced evenly spaced button presses that matched these global dynamics then the magnitude of the timing error would fluctuate from peak to peak. That is, if participants just tapped at a regular isochronous rhythm then timing error would vary as a function of peak position because the stimulus itself is not isochronous. If, on the other hand, participants adjusted their responses according to the local variations in the stimulus—that is, the local peak to peak timing variations—then timing error should be relatively constant across the trial. (The logic of this analysis is shown in Figure 5).

**Figure 5.**
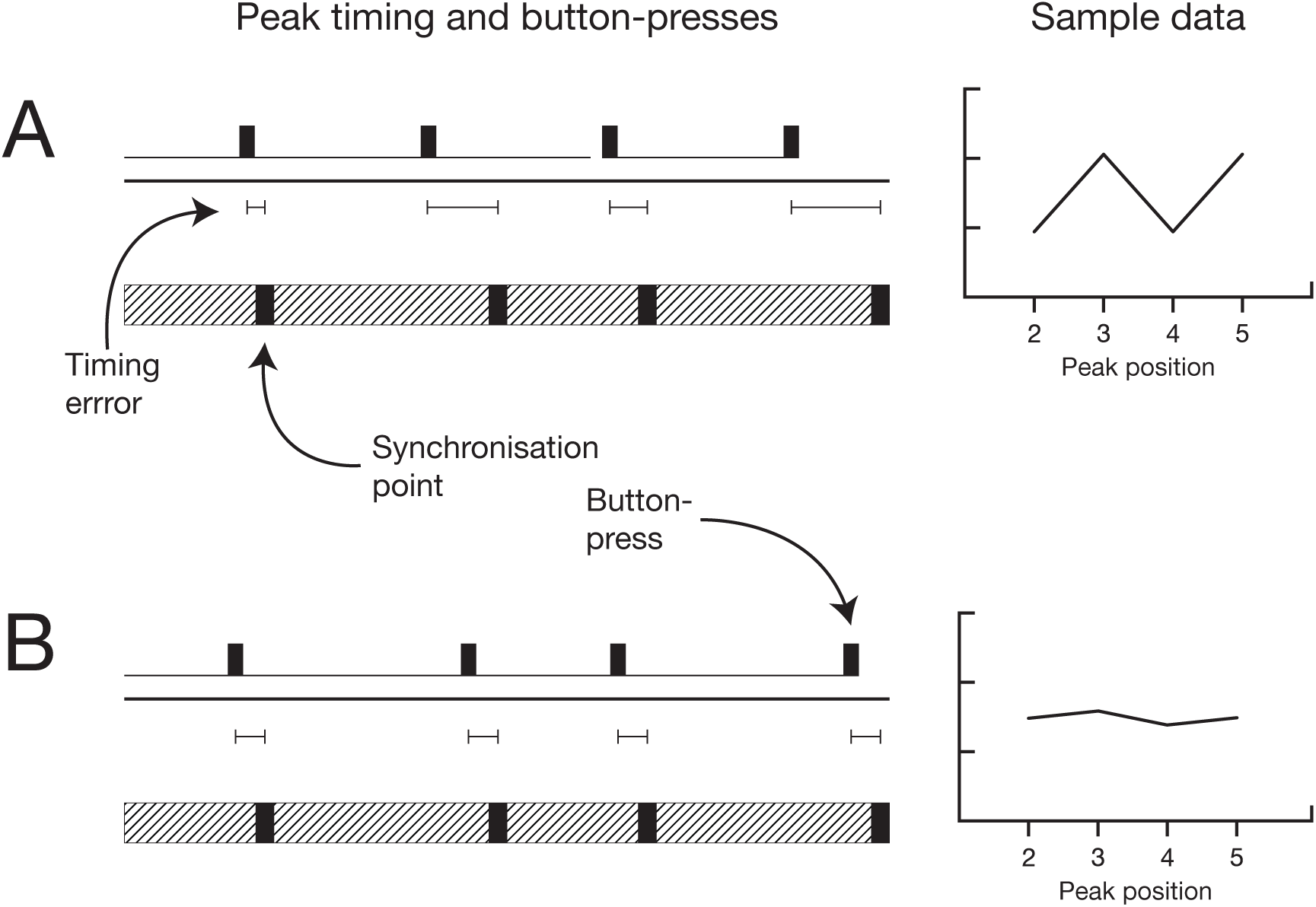
(A) Evenly spaced button-presses results in timing errors that vary as a function of peak number. (B) Button presses that vary according to peak position results in timing errors that do not vary according to peak position

In order to examine which of the two strategies was adopted by each of the groups, we analysed timing error using two separate one-way ANOVAs (with the factor Peak 2–5). If participants adopted the strategy of responding to the global dynamics of the stimulus, then this should be evident as a significant effect of peak position on timing error. However, if participants adopted the strategy of responding to the local dynamics of the stimulus then we should not find a significant effect of peak position of timing error. The results of the analysis showed a significant effect of peak position on timing error for the naïve group, *F*_3,33_ = 11.166, *p* = .005, ϵ = .374, 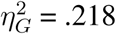, but not for the experienced group, *F*_3,36_ = 2.912, *p* = .105, ϵ = .396, 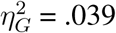. This is consistent with the experienced group and the naïve group adopting different strategies, with the naïve group responding to the global dynamics of the stimulus and the experienced group responding to the local dynamics. These data are shown in Figure 6.

**Figure 6.**
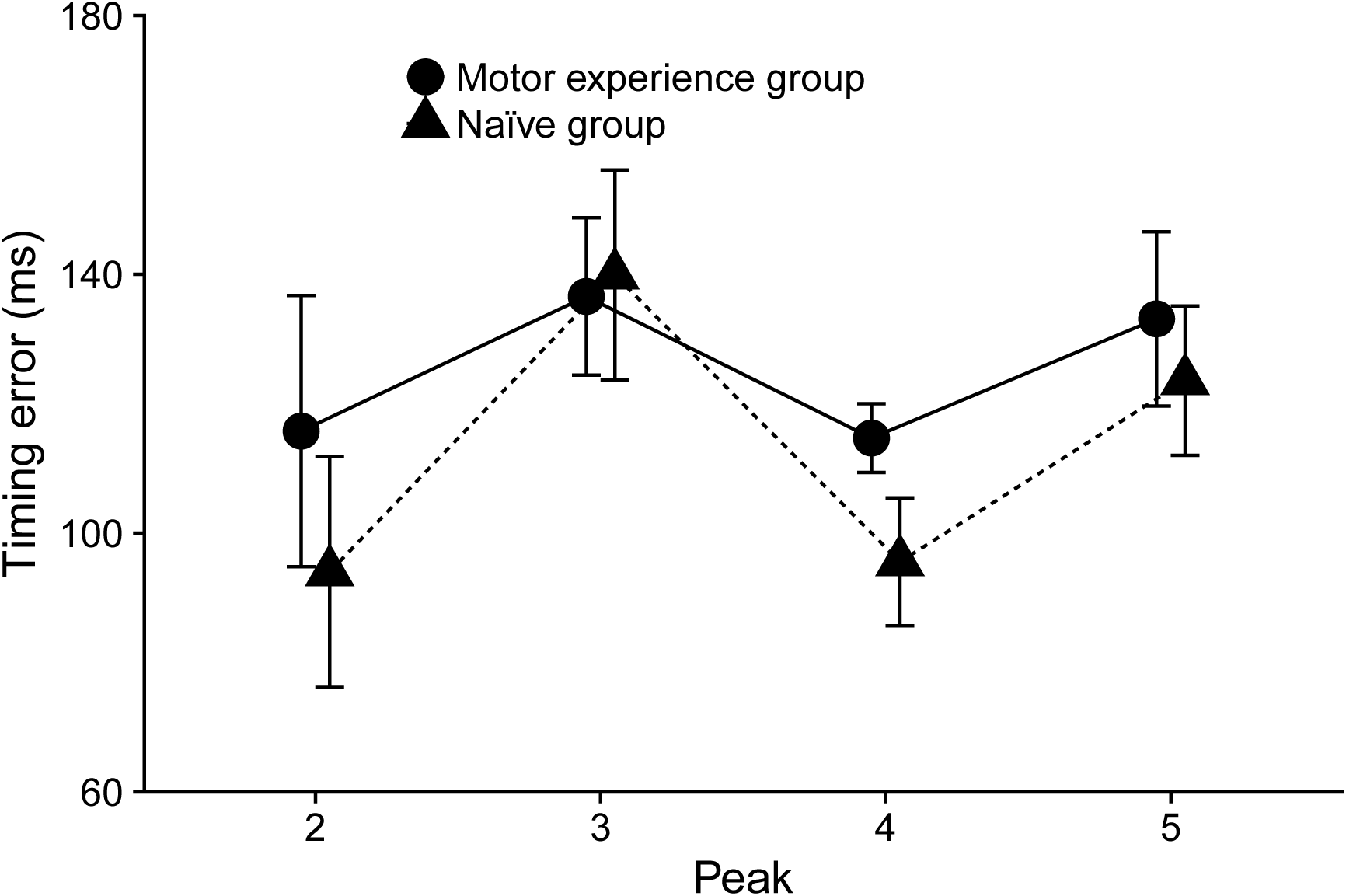
Timing error as a function of peak position for the experienced and naïve group. Error bars show the within-subjects 95% confidence intervals.

## Discussion

The primary aim of the present study was to investigate how online prediction of action timing is changed by motor experience. Previous studies have shown that observers who have experience performing an action are able to generate more accurate predictions about that action (e.g., Aglioti et al. 2008; Sebanz and Shiffrar 2009). This increase in prediction accuracy could be achieved in at least two ways. It might be that increased accuracy is achieved through motor experience fine-tuning or otherwise enhancing the operation of a predictive mechanism that is common to both naïve and experienced observers. However, it might also be the case that motor experience allows observers to engage different predictive mechanisms or apply distinct predictive strategies to the problem of action prediction, strategies or mechanisms that naïve observers are less able to call upon. By only measuring overall prediction accuracy, these studies only show *that* prediction is altered by motor experience but not *how* it is altered. Addressing this question was the aim of the present study. Specifically, we hypothesised that observers with motor experience would be capable employing predictive strategies that are action specific by using their knowledge of the specific actions being performed. Naïve observers, on the other hand, would only be able to engage a generic strategy as they do not possess any first-hand knowledge of the action.

We compared prediction accuracy for experienced and naïve participants under two stimulus conditions, which were designed to have a differential effect depending on whether the observer employed an action specific or generic strategy. If observers employed a non-specific strategy then altering the *action-related* properties of the stimulus—for example, the depiction of which effectors were used to produce the action—while holding the critical dynamic properties of the stimulus constant should have little or no effect on prediction accuracy, because the critical information—the stimulus dynamics—do not change between conditions. If, however, observers employed an action-specific strategy then changing *action-related* properties should have an effect on prediction accuracy. As hypothesised, we found that alignment accuracy was enhanced in response to the full information stimuli only for participants who had experience producing the observed action. Interestingly, we found no overall difference between the two groups that were independent of the condition and no overall difference between the conditions that was independent of the group.

While previous studies have been able to demonstrate *that* motor experience changes processes involved in action prediction by, for example, enhancing prediction accuracy (Aglioti et al. 2008), the results presented here go further to demonstrate *how* these predictive processes are changed. Specifically, these results are consistent with the idea that experienced and naïve participants rely on different mechanisms or strategies for action prediction. This distinction between internally replicating the action itself and merely simulating the stimulus dynamics within the motor system is similar to the distinction between *emulation* and *simulation*, respectively, put forward by Grush (2004). By internally replicating the action itself, observers might not only (in certain circumstances) generate more accurate predictions but may also generate predictions that more accurately replicate the fine-grained timing details of the observed action. These differences in fine-grained details may not appear in tests of gross performance, such as predicting binary action outcomes (e.g., Aglioti et al. 2008; Sebanz and Shiffrar 2009).

The findings of the present study are also consistent with recent TMS work by Novembre, Ticini, Schütz-Bosbach, and Keller (2014) and Hadley, Novembre, Keller, and Pickering (2015). Both these studies involved applying TMS over motor regions while participants’ were engaged in temporal coordination with stimuli that they either did or did not have experience with. For example, Novembre et al. (2014) had pianists play a duet along with a recording of a piece that they had also be trained to perform or with an untrained piece. The results showed the TMS was able to disrupt temporal coordination only when participants were playing along with a piece on which they had been trained. Similarly, in a musical turn-taking task, Hadley et al. (2015) found that TMS was able to disrupt the temporal precision of the participants’ entry into a joint performance only in trained but not untrained contexts. Taken together, these studies, as well as the results of the present study, show that temporal coordination with unfamiliar stimuli relies on different mechanisms or brain networks compared with temporal coordination with familiar stimuli.

Although not part of our initial hypothesis, we did conduct an exploratory secondary analysis to examine whether there was any information in the pattern of alignment accuracy data that would suggest that experienced and naïve observers were performing the task in different ways. In particular, if naïve participants generated their predictions using a generic, non-specific, simulation then we might expect these predictions to be less sensitive to fine-grained timing changes in the stimulus relative to the full-blown internal action replication that we hypothesised would be performed by the experienced observers.

To test this possibility, we compared the intra-trial differences in alignment accuracy for the two groups. The results showed that for the naïve group, alignment accuracy differed significantly as a function of peak position. This was not the case for the experienced group. This result could be produced by naïve participants merely responding to the global dynamics of the stimulus instead of responding to the fine-grained timing variations in the stimulus, as seen in the experienced participants. This result is consistent with the notion that experienced observers generate predictions about observed actions by employing an internal model of that action that is acquired through motor experience. By mapping the observed action onto their internal model for that action they are better able to capture the fine-grained timing variations in the stimulus because their predictive model more completely captures the constraints specific to the effectors used to produce the action. If naïve observers do not internally simulate the observed action then this generic model may be less capable of capturing these fine-grained details while still being able to capture the global dynamics.

Recent work using transcranial magnetic stimulation may also be relevant to the current work. Agosta, Battelli, and Casile (2016) examined cortico-spinal excitability during observation of action and non-action motion stimuli. While Agosta et al. (2016) found no difference in overall motor evoked potential (MEP) amplitude between the action and the non-action condition, with mean MEP amplitude only being sensitive to stimulus kinematics rather than stimulus form, differences in the temporal dynamics of cortico-spinal excitability were observed. In particular, it was found that the amplitude of the MEP correlated with the instantaneous velocity of the movement stimulus but not the abstract stimulus. This suggests that while non-action stimuli might, via mirror neurons, activate the motor system (consistent with the claims of Schubotz et al. 2004), this activation might be different in nature to the activation produced by action stimuli. Indeed, Agosta et al. (2016, p. 190) suggest that “observation of abstract motion [produce] a ‘coarser’ activation of the observer’s motor system”. This “coarser” activation, which less accurately tracks the fine-grained dynamics of the stimulus, might underlie the difference in prediction accuracy between the full information stimuli and point information stimuli reported in the present work. However, since Agosta et al. (2016) did not examine action prediction all that can be said is that their finding is consistent with the claims advanced here and not that they support our claims. An interesting avenue for future work, which may allow a further bridge to be built between the mirror neuron system and action prediction literature, would be to examine how the difference in the temporal dynamics of MEPs (reported by Agosta et al. 2016) are modulated by motor experience, perhaps using a task similar to the present or on an outcome prediction task such as that used by Aglioti et al. (2008).

We should also note that one might want to counter the claims presented by arguing that simulation or motor processes are not involved and that the effects observed here might instead be a result of observers getting better at predicting the percept. However, it’s not clear whether this is an alternative explanation to the one presented here. Rather, it seems that getting better at predicting the percept is the phenomena to be explained and the changes in the action prediction process outlined here is but one mechanism by which this can be achieved.

### The relationship between prediction strategy and expertise

It is important also that we not overstate our claims. Unlike studies by, for example, Calvo-Merino and colleagues (Calvo-Merino et al. 2006; Calvo-Merino et al. 2005) and Cross and colleagues (Cross et al. 2006), which suggest that motor experience might lead to the creation of new sensorimotor mappings, our claim is more modest. Rather than being the result of new sensorimotor mappings created by motor training, the effects observed here are instead more likely to indicate a switch in strategy—that is, a change in the flexible deployment of sensorimotor mappings. Knowledge about observed actions—that is, information about an action that is not directly accessible from the visual depiction—has been shown to influence effects such as, for example, motor priming. Motor priming, refers the phenomenon whereby observers are faster to produce an action when viewing a congruent action relative to an incongruent actionfor example, observers might be quicker to raise their index finger in response to a stimulus that also depicts a raised index finger relative to a stimulus that depicts a raised middle finger (e.g., see Brass, Bekkering, Wohlschläger, and Prinz 2000). One explanation for these effects is that observation of the action partially activates action production systems leading to quicker responses.

Importantly, however, these effects are not solely dependent on the action itself and can be modulated by giving the observer information about the action. For example, Liepelt, Cramon, and Brass (2008) demonstrated that whether motor priming occurs is dependent on the goal that observers attributed to the observed actors. The same physical stimulus could produce different effects, or different physical stimuli could produce the same effects, depending on the knowledge the observer possessed about the observed actions.

An analogous process might be responsible for the effects observed in the present study.

That is, rather than experience producing new sensorimotor mappings, the experience manipulation employed in our current study may change the way in which these sensorimotor mappings are flexibly deployed.

This raises the question of whether it would be possible to produce the effects observed here with some other kind of manipulation such as, for example, a simple verbal instruction of the type used in the motor priming literature. This remains a question for future research; however, we believe that there is likely to be a more complex relationship between the nature of the action itself and the manipulations that might lead to change in the strategy used to generate predictions about those actions. For example, with simple actions, such as finger movements, a verbal instruction might be sufficient. However, with more complex actions (salient) first-hand experience of actually performing the action might be necessary.

Furthermore, a switch in strategy might also depend on the visual representation of the action in a manner other than that explored in the present study. Although our study employed animated mannequins generated from motion capture data the mannequins were still far from human looking. It is likely to also be the case that using real humans (that is, videos) may change the way in which sensorimotor mappings are deployed and that these factors might also interact with experience and instruction-related factors. The work presented here provides only a preliminary glimpse into how action prediction processes might be modulated; however, we believe that this paradigm might be useful for addressing these questions in future research.

## Conclusions

Taken together, the results presented here suggest that observers with and without experience performing an action rely on different mechanisms or strategies when asked to generate predictions about that action. Observers who have experience actually performing the observed action may generate predictions by internally replicating the actual observed action, possibly through reactivating motor representations laid down by earlier performance. Observers without this experience, however, engage general purpose predictive mechanisms that do not necessarily replicate the actual action nor the fine-grained details of the observed action. Furthermore, when stimulus dynamics are held constant, only experienced observers are able to take advantage of action-related information (information about the limbs and joints used to produce the action) while this action-related information has no influence on the predictions generated by naïve observers. Thus, the findings of this study show not only *that* motor experience changes action prediction but also *how* motor experience changes the operation of these predictive processes. Furthermore, the results of the present study suggest the future work examining how experience modulates action prediction should, rather than employing a single task that cannot distinguish between different strategies for action prediction, employ manipulations that specifically tap into the predictive strategy of observers so that any differences in predictive strategy between experienced and naïve observers is evident.

## Acknowledgements

All the raw data, stimulus materials, analysis code, and supplementary figures are available online at http://osf.io/ebf45. We would like to thank Peter Keller, Harold Bekkering, Günther Knoblich, Natalie Sebanz, and Roman Liepelt, for their comments on an earlier version of this manuscript. We would also like to thank two anonymous reviewers for their comments.

## Footnotes

1 For neuroimaging studies that involve outcome prediction see, for example, Abreu et al. (2012) and Diersch et al. (2013).

2 See Keller, Knoblich, and Repp (2007) and Flach, Knoblich, and Prinz (2003) for similar tasks.

3 It is important to note that experience related changes in how sensorimotor mappings are employed during action prediction and expertise related changes in sensorimotor mappings should not be considered mutuality exclusive explanations. Rather, they operate at different parts of the action prediction mechanism.

4 The difference in age was statistically significant, t(12.33) = 2.42, p = .032. While age might influence overall performance, it is unlikely to account for the observed interaction.

5 An exploratory analysis found no significant effects or interactions with pattern type.

6 For more details on the prior distributions for each parameter, see Kruschke (2013).

